## Supplemental Tables for "Distinct Neuropeptide-Receptor Modules Regulate a Sex-Specific Behavioral Response to a Pheromone"

**Supplemental Table 1. Strains.**

| Strain | Genotype | Figure | Received From |
|---|---|---|---|
| N2 | wild-type, Bristol | Fig. 1, Supp. Fig. 1 | *Caenorhabditis* Genetics Center (CGC) |
| CB4088 | *him-5*(e1490) | Fig. 1, Supp. Fig. 1, 5 | CGC |
| CB1489 | *him-8*(e1489) | Fig. 1, 2, 3, 6, 7, Supp Fig. 1, 2, 3, 4, 6, 7, 9, 10 | CGC |
| PT690 | *osm-3*(mn391);*him-5*(e1490) | Fig. 1, Supp. Fig. 1 | Paul Sternberg (CalTech) |
| pk361 | *flp-3*(pk361) | n/a | Chris Li (CUNY) |
| NY183 | *flp-6*(pk1593) | n/a | Chris Li (CUNY) |
| NY106 | *flp-12*(n4902) | n/a | Chris Li (CUNY) |
| NY193 | *flp-19*(pk1594) | n/a | Chris Li (CUNY) |
| JSR99 | *flp-3*(pk361);*him-8*(e1489) | Fig. 2, 3, 6, 7, Supp Fig. 2, 3, 4, 5, 7, 9, 10 | This Study |
| JSR1 | *flp-6*(pk1593);*him-8*(e1489) | Fig. 2, Supp. Fig. 2 | This Study |
| JSR4 | *flp-12*(n4902);*him-8*(e1489) | Fig. 2, Supp. Fig. 2 | This Study |
| JSR2 | *flp-19*(pk1594);*him-8*(e1489) | Fig. 2, Supp. Fig. 2 | This Study |
| VC2497 | *flp-3*(ok3265) | n/a | CGC |
| JSR84 | *flp-3*(ok3265); *him-8*(e1489) | Supp. Fig. 3 | This Study |
| JSR109 | *him-8*(e1489);worEx17[p*flp-3*::*flp-3*::GFP; p*unc-122*::RFP];*flp-3*(pk361) | Fig. 2, Supp. Fig. 2 | This Study |
| JSR81 | *him-8*(e1489);worEx17[pflp-3::flp-3::SL2::mCherry (25 ng/uL); punc-122::GFP (50 ng/uL)] | Fig. 2, Supp. Fig. 2 | This Study |
| PS2218 | *dpy-20*(e1362);*him-5*(e1490);syls33[HS.C3(50ng/uL) + pMH86(11ng/uL)] | n/a | Paul Sternberg (CalTech) |
| JSR113 | *dpy-20*(e1362);*him-5*(e1490);syls33[HS.C3(50ng/uL) + pMH86(11ng/uL)];worEx27[p*flp-3*::*flp-3*::SL2::mCherry (25 ng/uL); p*unc-122*::GFP (50 ng/uL)] | Fig. 2 | This Study |

| | | | |
|---|---|---|---|
| tm1782 | *npr-4*(tm1782)) | n/a | National BioResource Project (NBRP) |
| CX14393 | *npr-5*(ok1583) | n/a | CGC |
| tm8982 | *npr-10*(tm8982) | n/a | NBRP |
| VC4220 | *frpr-16*(gk5305[loxP + p*myo-2*::GFP::*unc-54* 3' UTR + p*rps-27*::neoR::*unc-54* 3' UTR + loxP]) | n/a | This Study |
| JSR91 | *npr-4*(tm1782)*;him-8*(e1489) | Fig. 3, Supp. Fig. 7 | This Study |
| JSR97 | *npr-5*(ok1583)*;him-8*(e1849) | Fig. 3, Supp. Fig. 7 | This Study |
| JSR102 | *npr-10*(tm8982);*him-8*(e1489) | Fig. 3, Supp. Fig. 7 | This Study |
| JSR103 | *frpr-16*(gk5305[loxP + p*myo-2*::GFP::*unc-54* 3' UTR + p*rps-27*::neoR::*unc-54* 3' UTR + loxP])*;him-8*(e1489) | Fig. 3, Supp. Fig. 7 | Vancouver Node of International *C. elegans* Knockout Consortium |
| JSR107 | *npr-10(tm8982);frpr-16*(gk5305[loxP + p*myo-2*::GFP::*unc-54* 3' UTR + p*rps-27*::neoR::*unc-54* 3' UTR + loxP])*;him-8*(e1489) | Fig. 3, Supp. Fig. 7 | This Study |
| JSR126 | *npr-10*(tm8982);*him-8*(e1489)*;*worEx37[p*npr-10*::*npr-10*::GFP (25 ng/uL); p*unc-122*::RFP (50 ng/uL)] | Fig. 6, Supp. Fig. 9 | This Study |
| JSR133 | *frpr-16*(gk5305[loxP + p*myo-2*::GFP::unc-54 3' UTR + p*rps-27*::neoR::*unc-54* 3' UTR + loxP]);*him-8*(e1489);worEx41[p*frpr-16*::frpr-16::SL2::mCherry (25 ng/uL); p*unc-122*::GFP (50 ng/uL)] | Fig. 6, Supp. Fig. 9 | This Study |
| CU5248 | *ceh-30*(tm272);*him-5*(e1490);smIs26[p*pkd-2*::GFP] | Supp. Fig. 5 | Ding Xue (UC Boulder) |
| JSR98 | *flp-3*(pk361);*ceh-30*(tm272);*him-5*(e1490);smIs26[p*pkd-2*::GFP] | Supp. Fig. 5 | This Study |
| PT315 | *egl-3*(n150);*him-8*(e1489) | Supp. Fig. 6 | Maureen Barr (Rutgers) |
| PT440 | *aex-5*(sa23);*him-8*(e1489) | Supp. Fig. 6 | Maureen Barr (Rutgers) |
| PT436 | *bli-4*(e937);*him-8*(e1489) | Supp. Fig. 6 | Maureen Barr (Rutgers) |

**Supplemental Table 2. Plasmids**

| Plasmid ID | Function | Component | Array ID | Received From |
|---|---|---|---|---|
| pDONR p1-p2 | DONR Vector | L1-L2 sites | | |
| pL4440 | | Scramble Sequence | | Victor Ambors, UMass Medical School |
| JSR#DKR20 | ENTRY Clone | 6xHis-MRFGKR-SCRAMBLE_KRK_STOP | | |
| JSR#DKR27 | ENTRY Clone | 6xHis-MRFGKR-FLP3.1-KRK-STOP | | |
| JSR#DKR22 | ENTRY Clone | 6xHis-MRFGKR-FLP3.2-KRK-STOP | | |
| JSR#DKR12 | ENTRY Clone | 6xHis-MRFGKR-FLP3.4-KRK-STOP | | |
| JSR#DKR13 | ENTRY Clone | 6xHis-MRFGKR-FLP3.9-KRK-STOP | | |
| JSR#DKR23 | ENTRY Clone | 6xHis-MRFGKR-FLP3.10-KRK-STOP | | |
| pDEST-527 | DEST Vector | *E. coli* Expression Vector | | Dominic Esposito (Addgene plasmid # 11518) |
| JSR#DKR21 | *E. coli* Expession | 6xHis-MRFGKR-SCRAMBLE_KRK_STOP | | |
| JSR#DKR28 | *E. coli* Expession | 6xHis-MRFGKR-FLP3.1-KRK-STOP | | |
| JSR#DKR25 | *E. coli* Expession | 6xHis-MRFGKR-FLP3.2-KRK-STOP | | |
| JSR#DKR15 | *E. coli* Expession | 6xHis-MRFGKR-FLP3.4-KRK-STOP | | |
| JSR#DKR16 | *E. coli* Expession | 6xHis-MRFGKR-FLP3.9-KRK-STOP | | |
| JSR#DKR26 | *E. coli* Expession | 6xHis-MRFGKR-FLP3.10-KRK-STOP | | |
| JSR#DKR18 | Translational Fusion | p*flp-3*::*flp-3*::GFP | worEx17 | |
| JSR#DKR34 | Translational Fusion | p*flp-3*::*flp-3*::SL2::mCherry | worEx27 | GeneWiz |
| JSR#DKR35 | Translational Fusion | p*frpr-16*::*frpr-16*::SL2::mCherry | worEx41 | |

| | | | | |
|---|---|---|---|---|
| - | Co-Injection Marker | p*unc-122*::GFP | | Mark Alkema, UMass Medical School |
| - | Co-Injection Marker | p*unc-122*::RFP | | Shreekanth Chalasani, Salk Institute |
| DACR1432 | DONR Vector | L2-L3 SL2::dsRed::*unc-54* 3' UTR | | Josh Hawk, Yale University |
| pPD95.75 | GFP Fire Vector | GFP | | |

**Supplemental Table 3. Primers**

| Primer/ Ultramer Name | Primer/Ultramer Sequence | References | Final Strain |
|---|---|---|---|
| p*flp-3* Forward | GACTCTGCAGcatttccaagacacatttgacg | | JSR81, JSR95 |
| *flp-3* Reverse | GACTGGATCCttttccaaagcgcatggtt | | JSR81, JSR95 |
| p*npr-10* Forward | gtttgttttccggcactttc | | |
| *npr-10* Reverse_overhang | AGTCGACCTGCAGGCATGCAAGCTaattgaacttggaatcgtggtagt | | |
| p*npr-10* Forward#2 | cggcactttcctcatttttc | | JSR126, JSR137 |
| p*frpr-16* Forward | accgatttctgatcgacgtg | | |
| *frpr-16* Reverse_overhang | CAGCAGTTTCCCTGAATTAAAATTAaacaattgccggagcttttc | | |
| pfrpr-16 Forward#2 | ttctgatcgacgtgttggtt | | JSR133 |
| GFP_C | AGCTTGCATGCCTGCAGGTCGACT | Boulin et al. | |
| GFP_D | AAGGGCCCGTACGGCCGACTAGTAG | Boulin et al. | |
| GFP/dsRed_D#2 | GGAAACAGTTATGTTTGGTATATTGGG | Boulin et al. | |
| dsRed_E | TAATTTTAATTCAGGGAAACTGCTG | | |
| dsRed_F | AAAGTTGgaaacagttatgtttgg | | |
| SCRAMBLE Forward | GGGGACAAGTTTGTACAAAAAAGCAGGCTGGATGCGCTTTGGAAAACGTaattcgaagctccaccgcggtggcggccgctctagaactagtggatccaccggttccatggctagccacgcgcgtggatcccccgggctgcaggAAACGTaaataaCACCCAGCTTTCTTGTACAAAGTGGTCCCC | | |
| FLP-3.1 Forward | GGGGACAAGTTTGTACAAAAAAGCAGGCTGGATGCGCTTTGGAAAACGTtctccactgggaacaatgcgctttggcAAACGTaaataaCACCCAGCTTTCTTGTACAAAGTGGTCCCC | | |
| FLP-3.2 Forward | GGGGACAAGTTTGTACAAAAAAGCAGGCTGGATGCGCTTTGGAAAACGTactccattgggaactatgcgttttggaAAACGTaaataaCACCCAGCTTTCTTGTACAAAGTGGTCCCC | | |
| FLP-3.4 Forward | GGGGACAAGTTTGTACAAAAAAGCAGGCTGGATGCGCTTTGGAAAACGTaaccctcttggaaccatgcgctttggaAAACGTaaataaCACCCAGCTTTCTTGTACAAAGTGGTCCCC | | |

| | | | |
|---|---|---|---|
| FLP-3.9 Forward | GGGGACAAGTTTGTACAAAAAAGCAGGCTGGATGCGCTTTGGAAAACGTaatcctgagaacgacacaccattcggaacaatgagatttggaAAACGTaaataaCACCCAGCTTTCTTGTACAAAGTGGTCCCC | | |
| FLP-3.10 Forward | GGGGACAAGTTTGTACAAAAAAGCAGGCTGGATGCGCTTTGGAAAACGTtctactgttgattcttcggagcccgtcattcgtgatcagAAACGTaaataaCACCCAGCTTTCTTGTACAAAGTGGTCCCC | | |
| frpr-16-1 SeqVal R | ggtatctgtggtttatgtcggatag | | JSR103, JSR133 |
| frpr-16-2 SeqVal R | gaacggcatacgttcaggata | | JSR103, JSR133 |
| frpr-16-2 WT F | aaactaccctgtgcgattagg | | JSR103, JSR133 |
| frpr-16-1 WT R | CTTCCATACGTCAAGCACTTTC | | JSR103, JSR133 |
| frpr-16-1 SeqVal F | actttcaggcgaacacatact | | JSR103, JSR133 |
| pMYO-2 SEC Insertion | ccctcaatgtctctacttgt | | JSR103, JSR133 |
| NeoR-SEC Insertion | TTCCTCGTGCTTTACGGTATCG | | JSR103, JSR133 |
