## Supplemental Information for "Distinct Neuropeptide-Receptor Modules Regulate a Sex-Specific Behavioral Response to a Pheromone"

### **Supplemental Figure and Table Legends**

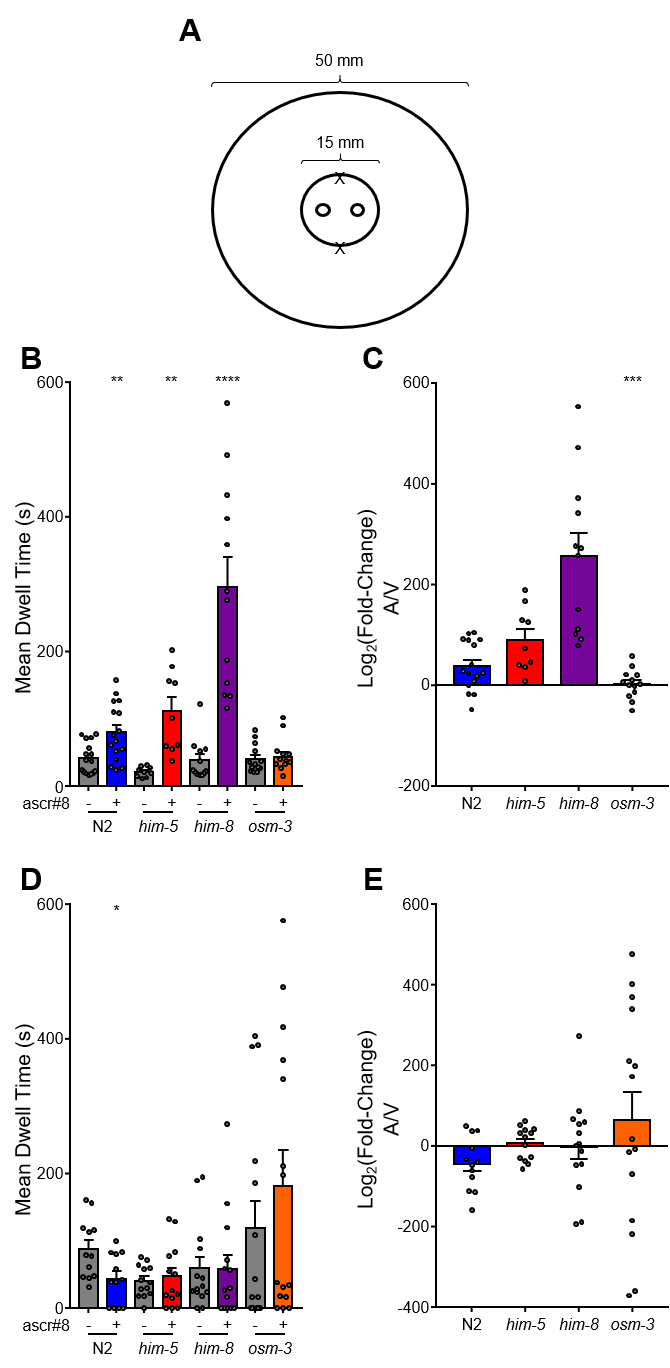
**Supplemental Figure 1. Canonical Spot Retention Assay. (A)** The Spot Retention Assay (SRA), as described previously ^14,33^. **(B)** Male *C. elegans* are attracted to ascr#8 in all wild-type strains (N2, *him-5*, *him-8*), but not chemosensory mutants (*osm-3;him-5*). **(C)** Transformed attraction data, logbase2(fold-change) ^38^ , of the raw data shown in **(B)**. **(D)** Hermaphroditic responses to ascr#8 across strains. **(E)** Transformed logbase2(fold-change) data of hermaphrodite SRA data.

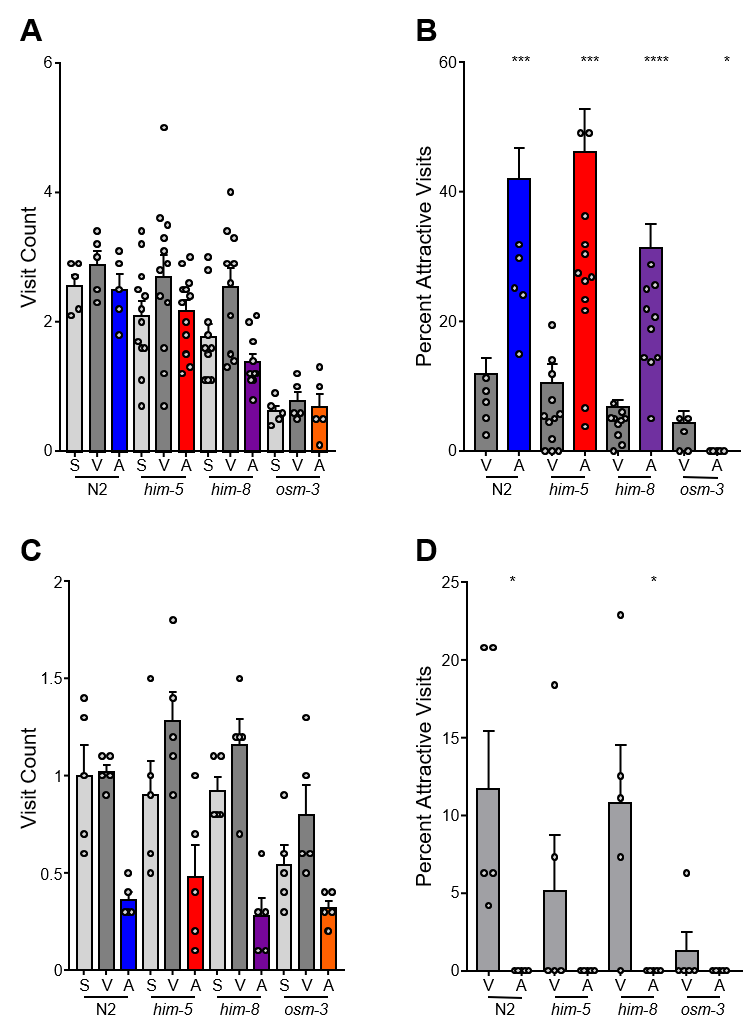
**Supplemental Figure 2. Single Worm Attraction Assay of control strain visit count and percent attractiveness. (A)** Male visit count of control strains. **(B)** Percent of attractive visits per worm for males of control strains. **(C)** Hermaphrodite visit count of control strains. **(D)** Percent of attractive visits per worm of control strains.

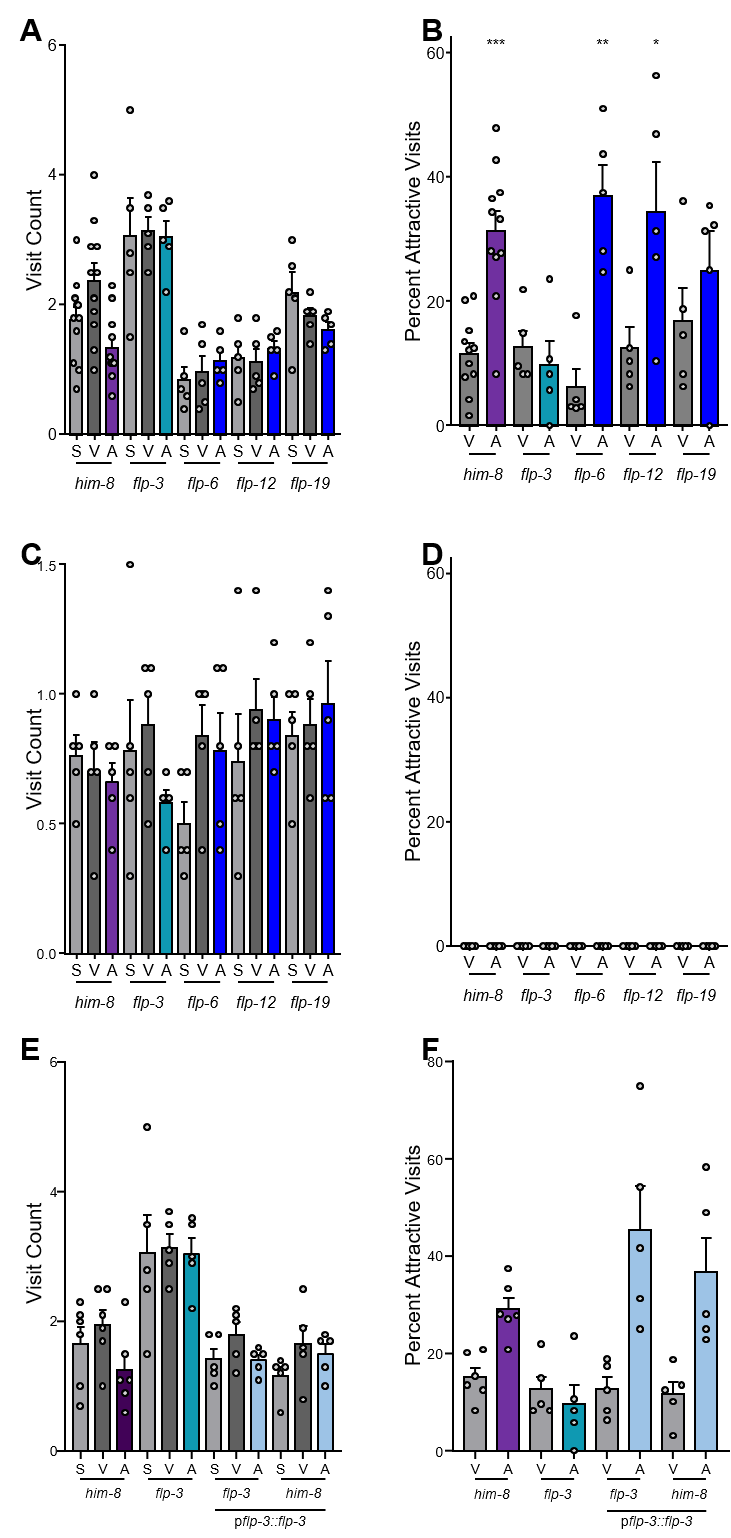
**Supplemental Figure 3. Neuropeptide null mutant screen SWAA supplemental information. (A)** Visit counts for males defective in neuropeptide genes. **(B)** Percent of attractive visits per worm of males defective in neuropeptide genes. **(C)** Hermaphrodite visit counts. **(D)** Percent of hermaphrodite attractive visits per worm. **(E)** Visit counts for males of *flp-3* rescue and overexpression transgenics. **(F)** Percent of attractive visits per male worm of *flp-3* rescue and overexpression transgenics.

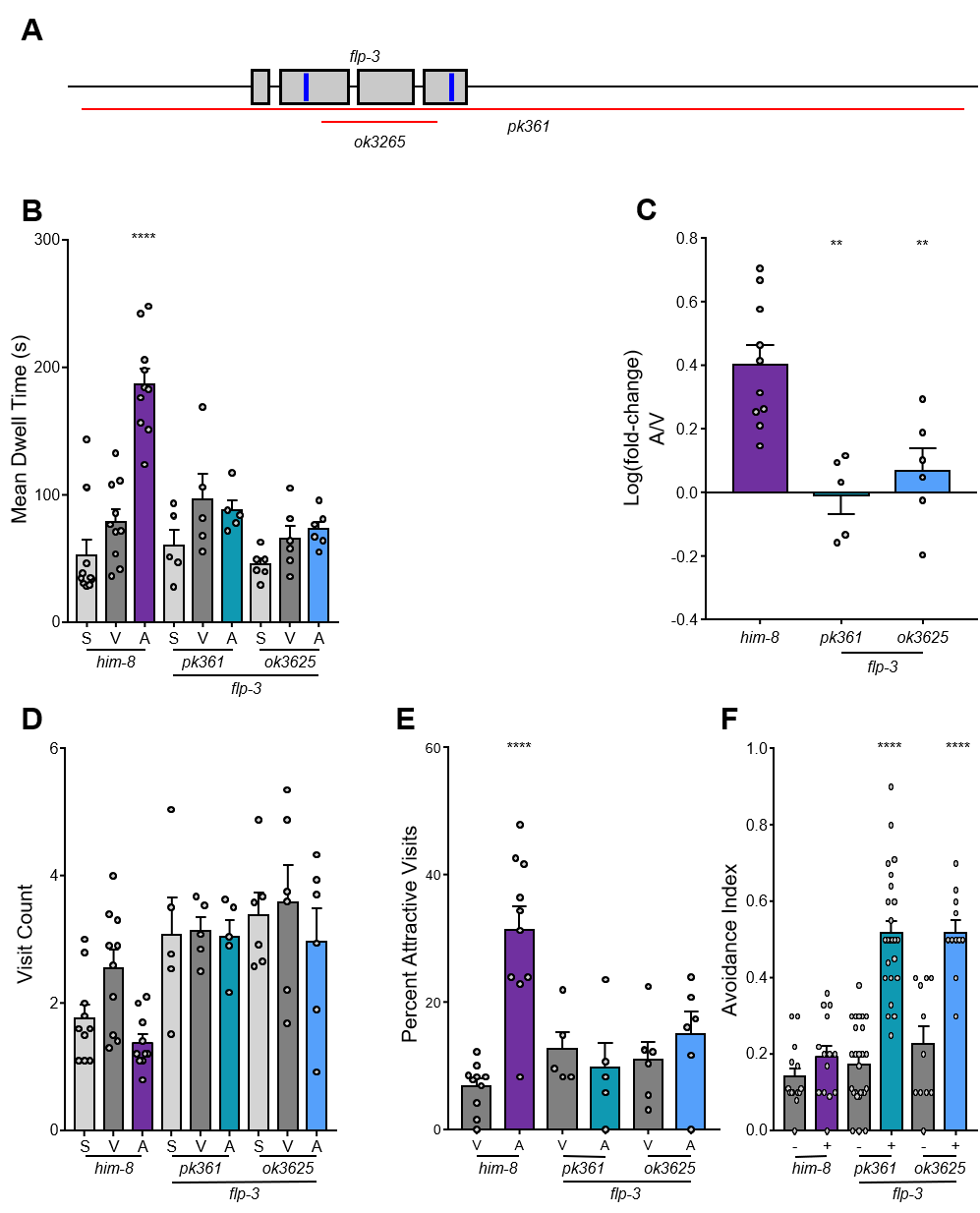
**Supplemental Figure 4. The *flp-3* phenotype is consistent across alleles. (A)** Schematic of *flp-3* deletion alleles *pk361* and *ok3625*, which result in a full gene deletion and in-frame partial gene deletion, respectively (red lines). **(B)** Raw dwell time, **(C)** log(fold-change) values, **(D)** visit count, and **(E)** percent of attractive visits of both *flp-3* alleles, *pk361* and *ok3625*. **(F)** Avoidance indexes of both *flp-3* alleles.

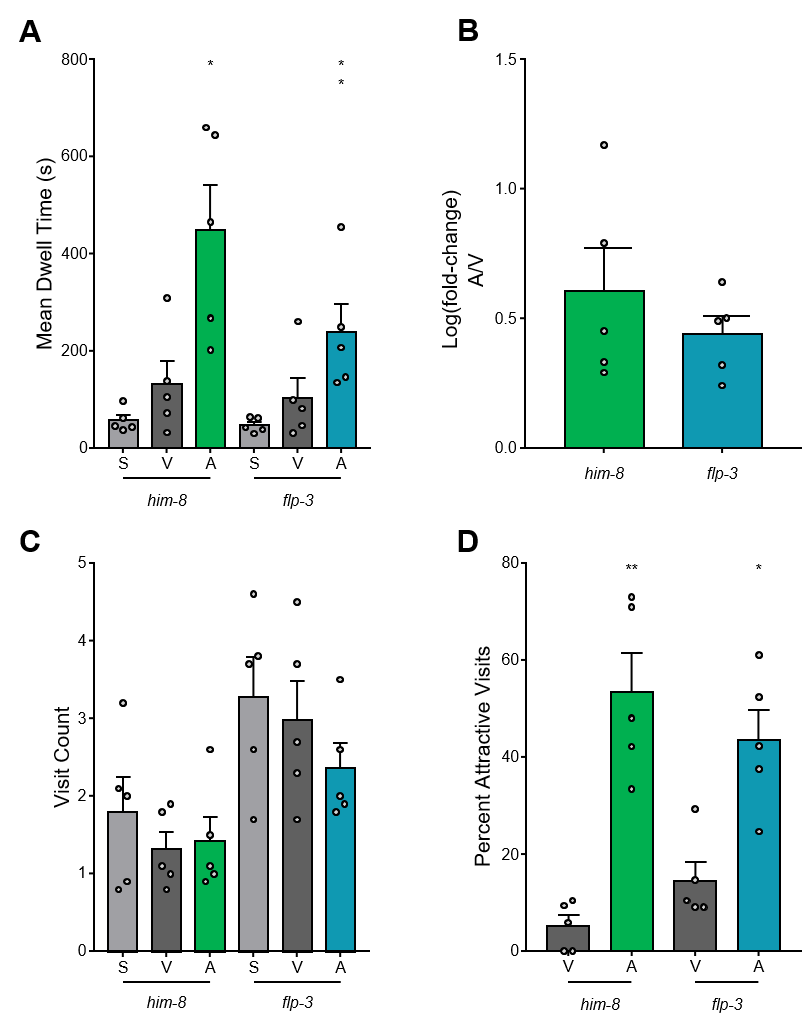
**Supplemental Figure 5. The Role of *flp-3* is Specific to ascr#8. (A)** Raw dwell time, **(B)** log(fold-change) values, **(C)** visit count, and **(D)** percent of attractive visits of *flp-3(pk361)* to two attractive ascarosides, ascr#8 and ascr#3.

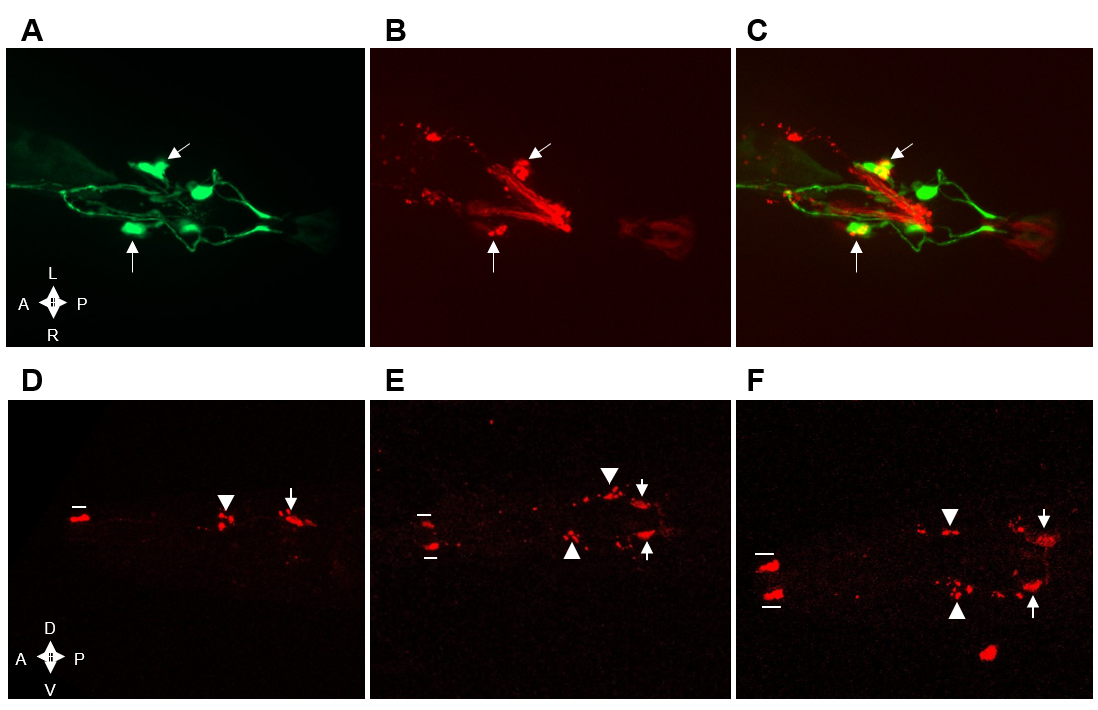
**Supplemental Figure 6. Expression pattern analysis of p*flp-3*::*flp-3*::mCherry. (A-C)** ~90x magnification of a mail tail expressing **(A)** *gpa-1*::GFP and **(B)** p*flp-3*::*flp-3*::mCherry. **(C)** The two reporters co-localize in the SPD neuronal soma (arrows). **(D-F)** IL1 expression of p*flp-3*::*flp-3*::mCherry. mCherry is faintly observed in the IL1 soma (arrows). The fluorescent protein is also observed in the dendritic cilia of the IL1 neurons (bars), as well as in punctate vesicles along the dendrites (arrowheads).

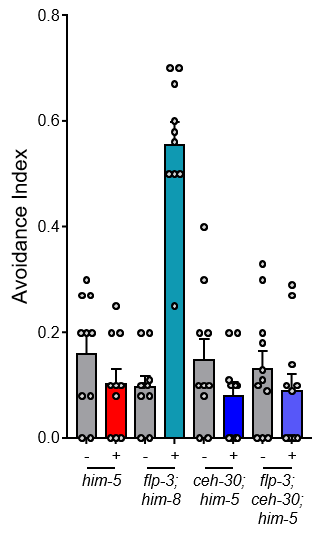
**Supplemental Figure 7. The CEM neurons are the sole chemosensory pathway detecting ascr#8 in males.** Avoidance index values of *him-5* and *flp-3;him-8* control males, as well as *ceh-30;him-5* and *ceh-30;flp-3;him-5* males.

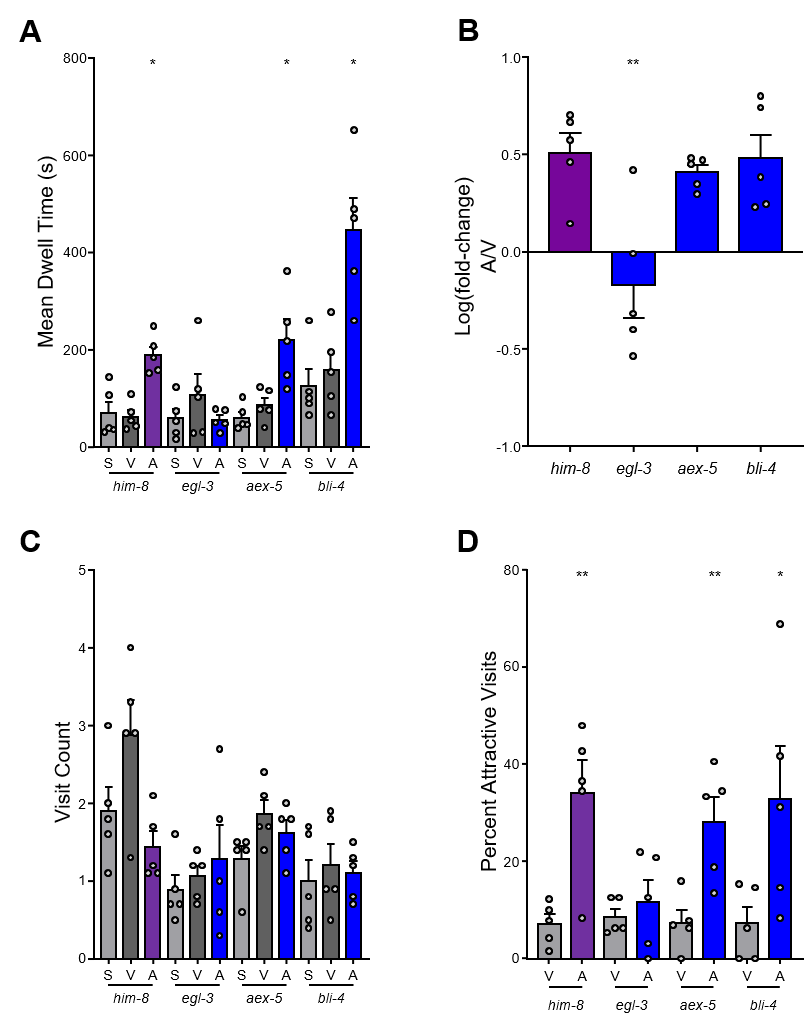
**Supplemental Figure 8. EGL-3 is the proprotein cleavage enzyme responsible for FLP-3 maturation. (A)** Raw dwell time, **(B)** log(fold-change) values, **(C)** visit count, and **(D)** percent of attractive visits per worm of propeptide convertase enzymes involved in neuropeptide maturation: *egl-3*, *aex-5*, and *bli-4*.

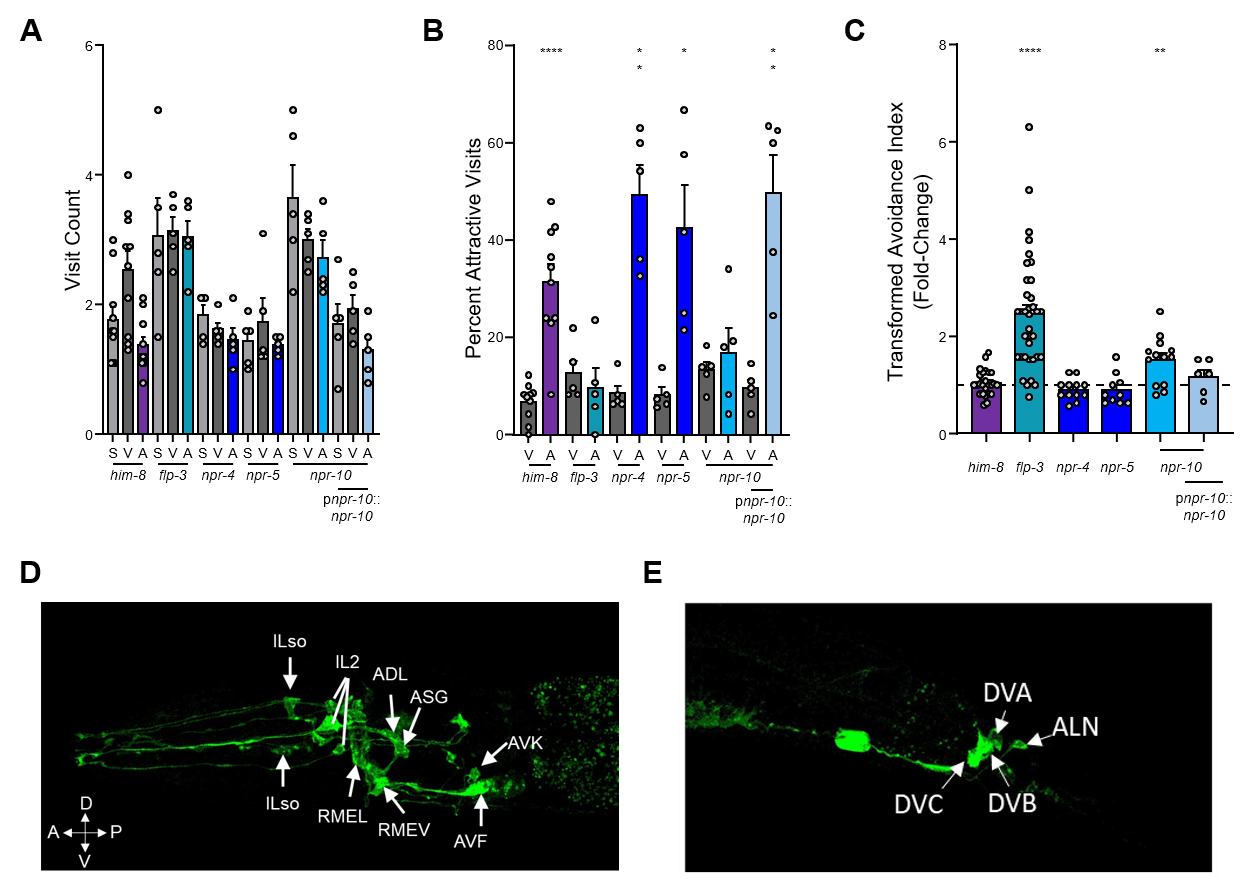
**Supplemental Figure 9. NPR SWAA supplemental information. (A)** Visit count and **(B)** percent of attractive visits per worm of *npr* receptor mutants and *npr-10* rescue. **(C)** Transformed avoidance of *npr* mutants and *npr-10* rescue. **(D)** Amphid localization of *npr-10*::GFP in the head of a hermaphrodite. **(E)** Localization of *npr-10*::GFP in the tail of the hermaphrodite, with expression observed in the dorsal-rectal ganglia neurons: DVA, DVB, DVC, and ALN.

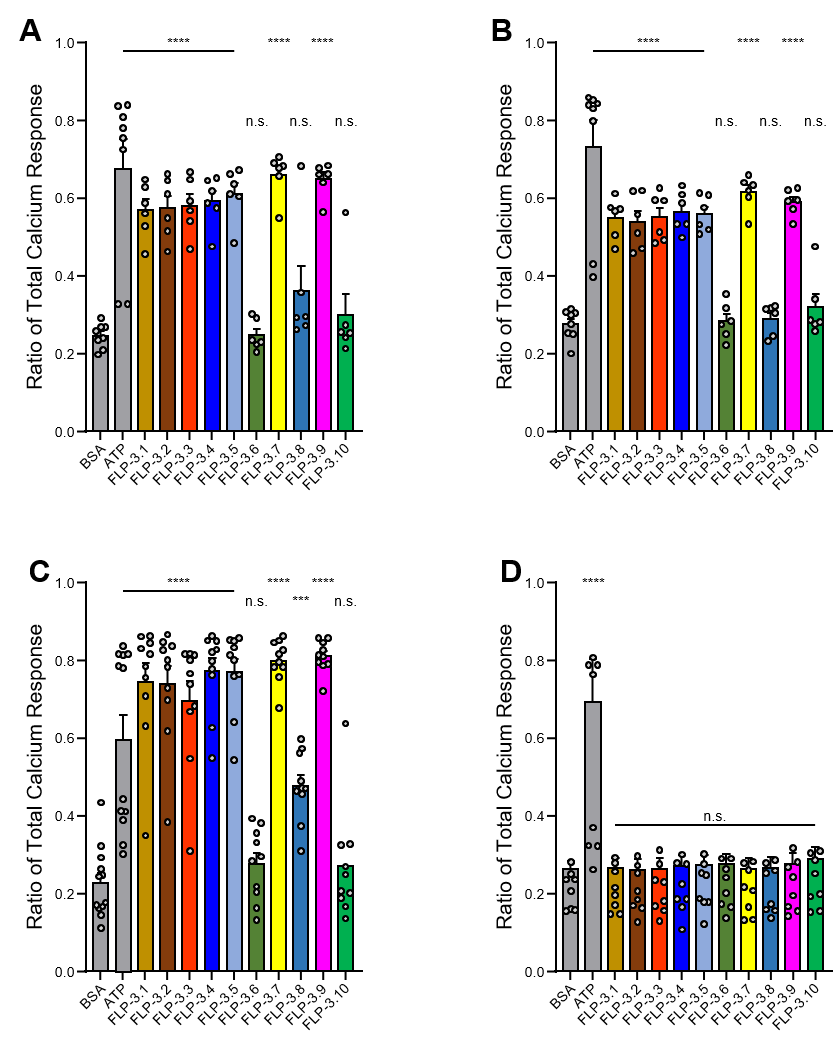
**Supplemental Figure 10. FLP-3 Peptides Activate NPR-10 and FRPR-16 *in vitro*.**

Ratios of Total Calcium Response for cells transfected with **(A)** NPR-10A, **(B)** NPR-10B, **(C)** FRPR-16, and **(D)** empty vector control. Peptides FLP-3-6 and FLP-3-10 activated no receptors, likely due to lack of sequence homology. No peptides activated cells transfected with empty vector control.

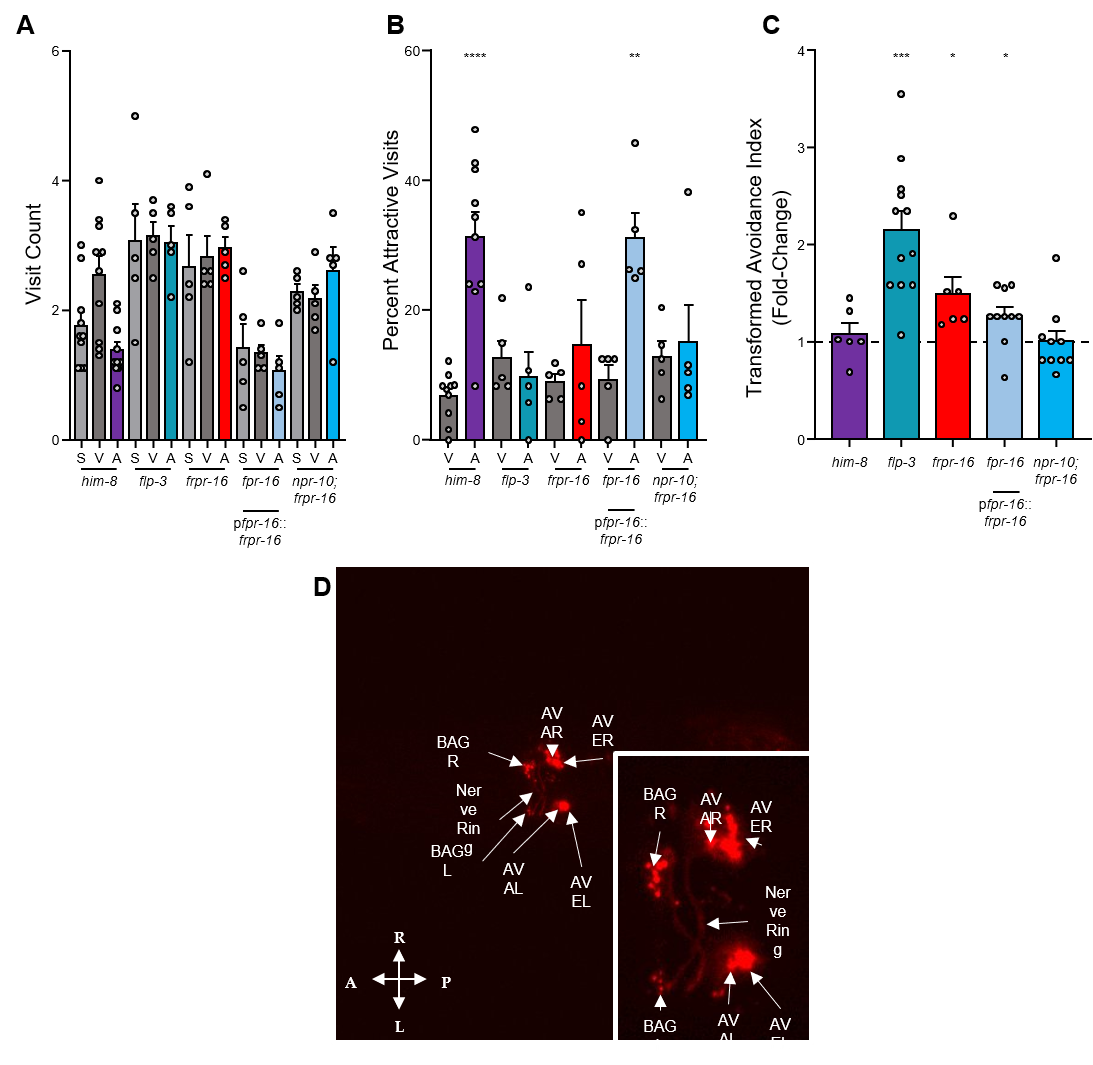
**Supplemental Figure 11. FRPR-16 supplemental information. (A)** Visit count and **(B)** percent of attractive visits per worm of *frpr-16 lof* animals, transgenic rescues, and *frpr-16;npr-10* double mutant animals. **(C)** Transformed avoidance values for *frpr-16 lof* animals, transgenic rescues, and *frpr-16;npr-10* double mutant animals. **(D)** Amphid localization of p*frpr-16*::*frpr-16*::SL2::mCherry in the head of a hermaphrodite. Similar expression within the AVA, AVE, AVD, and BAG neurons is seen. **(D, inset)** 2X zoom of main image, focusing on nerve ring localization.

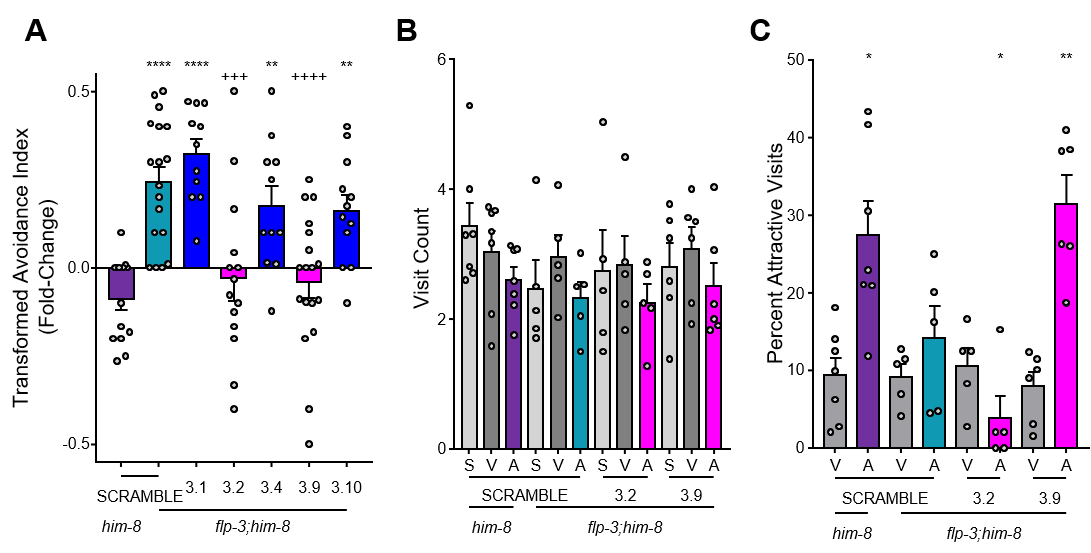
**Supplemental Figure 12. FLP-3 peptide feeding rescue supplemental information. (A)** Transformed avoidance of *flp-3* animals fed SCRAMBLE and FLP-3 peptides. **(B)** Visit counts, and **(C)** percent of attractive visits per worm of *flp-3* animals fed peptides.
